# The Effect of Amniotic Fluid On Haıir Follicle Growth

**DOI:** 10.1101/2023.09.08.556806

**Authors:** Gamze Tumentemur, Elif Ganime Aygun, Bulut Yurtsever, Didem Cakirsoy, Ercument Ovali

## Abstract

Human amniotic fluid stem cell (hAFSC) exhibit significantly as a new treatment modality in hair loss, wound, scar and regenerative plastic surgery/dermatology. The difference of our study from previous amniotic fluid studies on the hair growth model was studied through the direct effect of human amniotic fluid (hAF). In our study, amniotic fluid was made acellular, pooled and freezed, making it more standardized in terms of its contents. Therefore, the present study is aimed at investigating the efficacy and safety of amniotic fluid in hair loss model on rat. Using the hair loss model rats, we investigated the therapeutic potential of freezing amniotic fluid (FAF) and freezing gama irradiated amniotic fluid (FAFI). Our results showed that FAF and especially FAFI increased number of total hair follicles and accelerated number of transition to anagenic hair follicle. We observed that increased rate of arginase+1(Arg+1)/CD68 activity around the hair follicles treated with hAF. Our study suggests that FAFI may represent a safe and effective tool for increasing hair follicle and accelerating to anagen stage during hair loss model.

## 1. Introduction

Hair loss (alopecia) is a common birth, hormonal, traumatic and iatrogenic problem. Alopecia Areata (AA) is an autoimmune hair disorder [1] and androgenic alopecia (AGA) is a condition that occurs with the effect of androgens, can be seen progresses with hair loss in male genders. Oral and topical drug or surgical options for the treatment of alopecia areata. Topical drug treatment includes corticosteroids, minoxidil and immunotherapy [2]. Contact immunotherapy with diphenylcyclopropenone is mainly used for limited hair loss [3]. However, these drugs have inadequate treatment methods because of side effects and limited therapeutic uses [4]. Recently, the efficacy of many cytokine-targeted drugs has been described in patients with AA [5]. Further, autologous single follicle and follicular unit transplantation has been a reliable surgical option, but the number of donor follicles limits this method.

Hair follicle (HF) is a mini-organ containing epidermal and dermal layers that go through hair cycle processes to produce new hair continuously throughout the life of the organism [6]. Hair follicle stem cell (HFSC) niche is responsible for maintaining tissue homeostasis in response to physiological and pathological conditions. It contains many cell types, including intradermal adipocytes [7], dermal fibroblasts [6], cutaneous blood vessels [8], macrophages and endothelial cells [9]. Immune cells in the hair follicle epithelium are called the hair follicle immune system [10]. Macrophages and T lymphocytes are a cutaneous immune cells. Dermal papilla (DP) provides signals that control hair follicle development and contribute to determining the size, shape and pigmentation of the hair shaft [11;12] as well as acting as a reservoir of stem cells [13]. An altered the DP microenvironment can lead to human skin hair loss, such as androgenic or chemically induced hair loss [14; 15;16]. The hair cycle is divided into three phases: growth and regeneration, regression, resting phase (anagen, catagen and telogen, respectively). There is a connection between macrophages and the regulation of the HF cycle, particularly between anagen-catagen transition [10;17]. During the catagen phase, macrophages digest the excess extracellular matrix and stimulate follicular stem cells, thus stimulating HF to enter the anagen phase [18]. Macrophages entering the anagen stage perform collagen phagocytosis to remodelling the matrix [19].

Hair cycle and regeneration which are the tissue restructuring process that includes growth factors, cytokines, hormones, adhesion molecules and related enzymes [20]. Growth factors (Insulin like growth factor-(IGF-1), Platelet derived growth factor (PDGF) and fibroblast growth factor (FGF) play important roles in HF morphogenesis by acting on stem cells in the bulge area of hair follicles [21; 22]. Although positive effects of growth factors on hair follicle regeneration have been reported, their high cost limits using in clinical applications [23]. The use of stem cells and growth factors in cell-based therapies are accepted as a therapeutic strategy in damaged tissue repair due to their direct cellular effects [24]. But human amnion-derived stem cells (hADSCs), human amniotic epithelial stem cells (hAESCs) and human amniotic mesenchymal stem cells (hAMSCs), have great advantages compared to other stem cells such as regeneration, tumorigenicity former. Moreover, there is no ethical provision or legal restrictions as embryonic stem cells are obtained from an adult source [25;26]. Furthermore their high proliferation capacity, multipotency, immunomodulatory effects make them a promising source of stem cells for cell therapy in various diseases [27; 28; 29; 30; 31]. As medical wastes, AFSCs can be obtained via amniocentesis during the pregnancy intermediate stage or cesarean section [32]. Ability of AFSCs to multi-differentiate is limited, but they do not have the risk of teratoma formation [33].

Amniotic fluid (AF) is a mixed biological fluid containing protein, lipid, carbohydrates, enzymes, urea, electrolytes, hormones, growth factors (IGF-1, PDGF, IL-8, IL-6, transforming growth factor (TGF)-β, TNF-α, vascular endothelial growth factor (VEGF), and epidermal growth factor (EGF)), stem cell and stem cell exosomes [34; 35; 36; 37; 38; 39; 40; 41; 42]. AF and Amniotic fluid-derived mesenchymal stem cells (AF-MSCS) which are known to be important in normal wound healing, are thought to trigger cell proliferation, differentiation, angiogenesis and chemotaxis, which are necessary for new hair follicle growth [41]. AFSC and it’s cytokines especially macrophages involved in HF mesenchyme, have interesting regulatory properties in hair follicle homeostasis [17; 43; 44]. Disrupting between this connection may lead to clinically important forms of immune-mediated alopecia [45; 46; 47].

Studies of amniotic fluid on alopecia have been demonstrated through the stem cell effect of this fluid. Our study, amniotic fluid was made acellular and pooled, making it more standardized in terms of its contents. In terms of content, it has been made safe even against viral and pyrionic contamination [48] due to gamma radiation. Therefore, it was designed to investigate the effect on the rat hair follicle development model by creating a longer shelf life and more effective form.

## 2. Results

### 2.1 Amniotic fluid analysis results

As shown Fig. 2A, TGF-B (p<0.00714) and VEGF (p<0.00003) levels in FAFI group were higher than FAF group. When Th1 type cytokines were divided by the amounts of Th2 type cytokines, the values were found to be below one (Fig. 1B).

**Figure 1.**
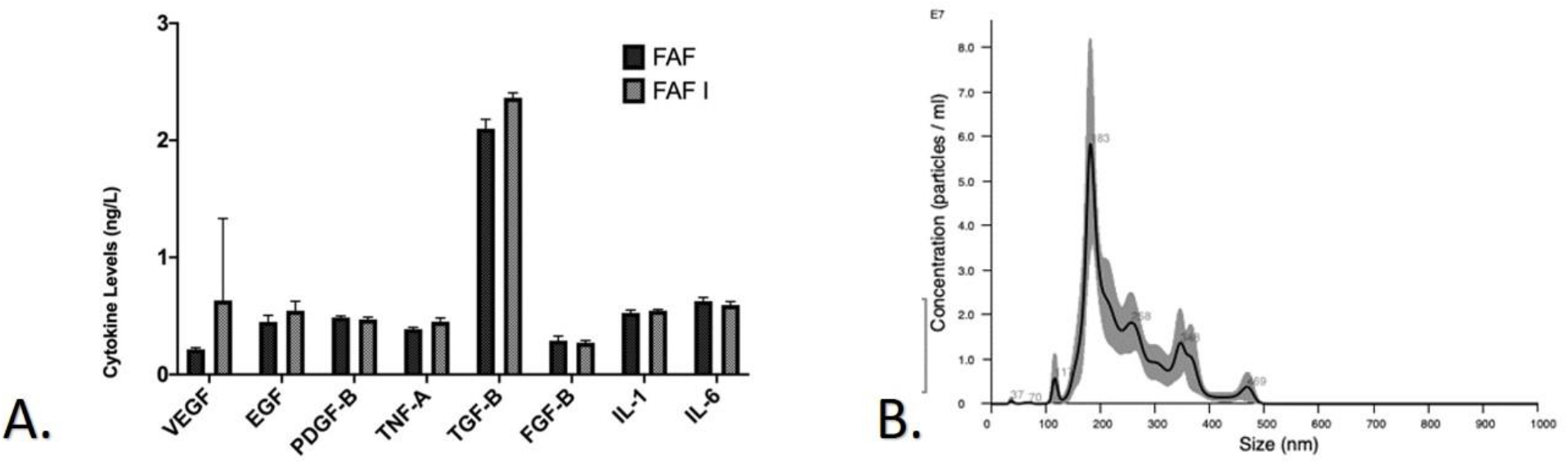
A statistical comparison of the cytokine levels of VEGF, EGF, PDGF-B, TNF-α, TGF-β, FGB-B, IL 1 and IL 6 levels in FAF and FAFI groups in an experimental hair follicle development model (ng/L) (A). Our data show that transition of M1 to M2 in treated with hAF groups that the amniotic fluid is contain high exosome content (1.14e10±0.531e10) (B).

**Figure 2.**
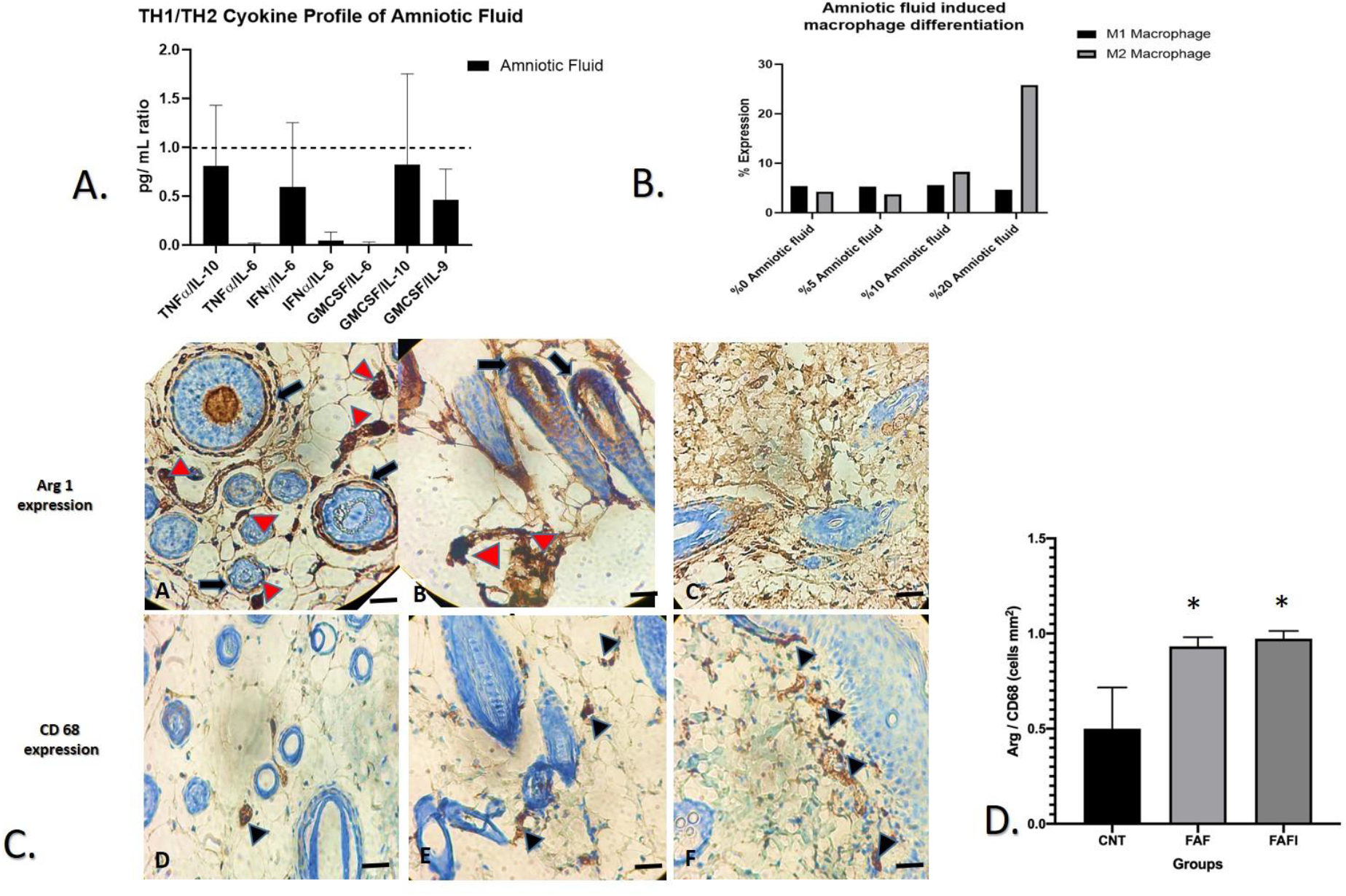
Differential hAF expression with macrophage differentiation (A). These data showed that amniotic fluid contains Th2 type cytokines (B). Macrophage cell counts in skin specimens (C). The figure shows the Arg+ and CD68+ cells count (cells per mm2 specimen area) in the whole analysed area in the stroma compartment of the skin specimens (D). The ANOVA test’s P-value results are shown.

### 2.2 Macrophage 2 Profile of Amniotic Fluid and Macrophage 2 Profile in Tissue

Our study showed that Th2 cytokines are linked to changes in expression levels of these cytokines (Fig.2 A). Our results that the Th2 type cytokines of the concentration of 10% and above amniotic fluid can induce the M1 to M2 phenotype switch in macrophages (Fig. 2B). The figure shows the expression of the M2 macrophage marker Arg+ cells around dermal papilla (black arrow) and stroma (red arrowhead) in samples (A:FAFI; B: FAF; C:CNT). The figure shows the expression of the CD68 +cells around dermal papilla (black arrowhead) in samples (D:FAFI; E: FAF; F:CNT) Scale Bars, 400 μm (Fig. 2C). A significantly increased rate of Arg+1/CD68 cells found in the FAF and FAFI specimens compared to cnt group (*p<0,05) All data are reported as the mean ± SD (Fig. 2D).

### 2.3 Macroscopic and Microscopic Analysis Results

#### 2.3.1 Effects of the FAF and FAFI on transition to anage phase in rat

Comparison of hair follicle pattern in each group on macroscopic evaluation histological observation to vertical view of hair follicles from the dorsal portion of rats were clearly visible (Fig. 3A-C). Compared to the cnt group, the transition to the anagen phase was accelerated (Fig. 3B-D) in the hAF-administered groups (Fig. 3D). At day 21, the hair follicle in the cnt group were in the early anagen phase as hair bulb in the dermis. The hair follicles in the FAF and FAFI groups, were in the late anagen phase and displayed the largest hair bulb size, the deepest hair follicle in the subcutis, and the newly formed hair shaft reaching the level directly below the sebaceous gland. Transition to anagen phase of hair follicle was quantified using histology images with Image J software (D-10×). All data are reported as the mean ± SD. *: p< 0,05; **: p<0,0001 compared with control,***: p<0,0001 compared with FAF group.

**Figure 3.**
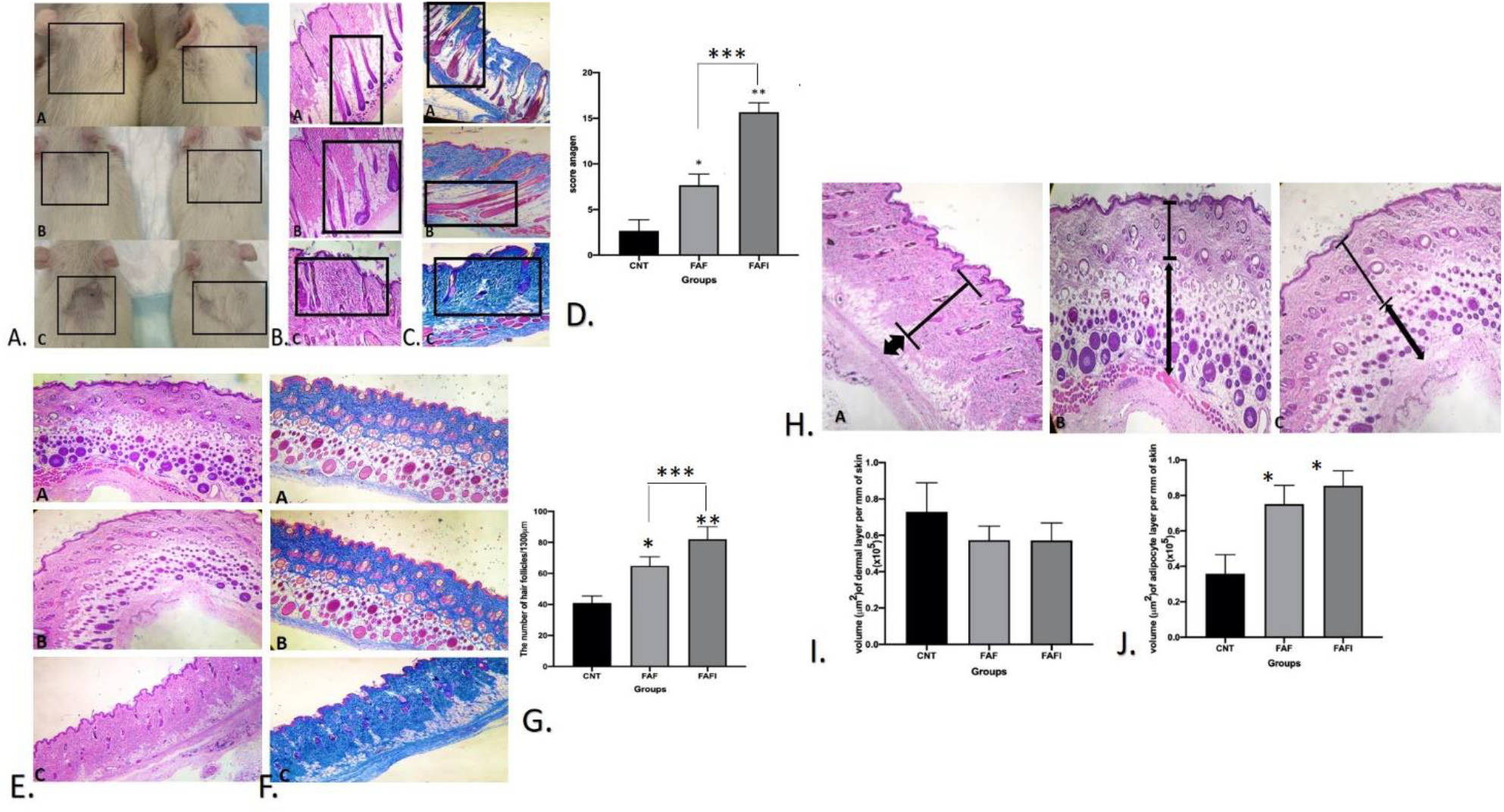
The black boxs show the hair follicle development on rat (A) and anagen stages in the sections (B-C). The rat hair follicle development was evaluated by H&E staining (B), masson trichrome (C) and anagen scoring (D). Typical photos of dorsal skin (left panel), histological analysis (right panel) (**A:** FAFI **B:** FAF, **C:** Cnt×100). Representative photomicrographs stained with H&E (E) and masson trichrome (F) of the transverse sections and hair follicle scoring (G). After 21 days, total hair follicle in the skin was analysed, number total hair follicles was quantified using histology images with image J software (C,10×). (**A:** FAFI, **B:** FAF and **C:** Cnt groups) ×100. p<0,001 **: p<0,0008 compared to cnt group ***: p<0,04 compared to FAF group. Relationship between the thickness of dermal layer and the thickness of the dermal adipocyte layer. Double-headed arrows indicate adipocyte layer, whereas bars indicate dermal layer (H). cnt (A), FAF (B), and FAFI (C) (H&E Scale bar, 200 μm). Quantification of the dermal (I) and the adipocyte layer thickness (J) in rat. Between the all groups, no different of dermal layers thickness layer (p˃0,05). Thickness of adipocyte layer that significantly increased FAF and FAFI groups compared to cnt group (*p<0,05) (J). Two-way ANOVA followed by Bonferroni’s multiple comparison test. All data are reported as the mean ± SD.

#### 2.3.2 Effects of the FAF and FAFI on hair growth in rat

We performed a histological evaluation hair follicles and collagen fibers using H&E and Masson staining respectively (Fig. 3E-F). In this model, more hair follicle regeneration was detected in the groups treated with hAF compared to the cnt group (Fig. 3E). Especially FAFI and FAF group exhibited a lower level of collagen fibers accumulation, which was more neatly arranged, compared with that in the cnt group (Fig. 3F). In the representative longitudinal sections, the number of hair follicles were significantly increased in the FAFI and FAF group compared to the cnt group (respectively p<0.001 p<0.0008) (Fig. 3G). In addition, the number of hair follicles in the FAFI group was significantly greater than FAF group (Fig. 3G).

#### 2.3.3 Effects of the FAF and FAFI on adipocyte and dermal layer in rat

We performed a histological evaluation of the thickness of dermal and adipocyte layer using H&E staining (Fig. 3H). We quantified the thickness of the dermal and adipocyte layer in dorsal rat skin on 21th day (Fig. 3I-J). There was no change the dermal layer between all the groups (p˃0.005) (Fig. 3I). We showed that the adipocyte layer increased in thickness during HF morphogenesis in the FAFI and FAF groups (p<0.005) (Fig. 3J).

## 3. Discussion

The present study provides the first report of hair growth and transition to anagen phase promotion by hAF whether FAF and FAFI groups can effective in promoting hair growth, were investigated. However, it is well known that stem cell of hAF has the activity on not only hair follicle regeneration but also transition of hair follicle from catagen to anagen [41; 51; 52; 53]. The present investigation well documented the direct effect of hAF on the hair growth. A striking finding in our study was that hAF promoted catagen-to-anagen transition. Our current histological analysis suggest that hAF was accelerated hair follicle regeneration and reduce collagen fiber deposition. Furthermore, the significant anagen transition and hair growth were seen in FAFI group. There are two points under discussion in terms of these findings. The first is the question of why irradiated group is more effective than non-irridated group. This previous study show that gamma irradiation can cause growth factors to be kept more stable by inhibiting the protease activity that causes the destruction of active substances in a liquid [48]. Our present finding was also related to this protective effect of gamma irradiation.

Treatment of hair loss using growth factors show interesting activity in promoting hair growth. Choi et al. reported the TGFβ family is very important for HF morphogenesis and hair cycle by triggering HF formation with DP activation [54]. In addition, TGF-β has been reported to increase Arg activity [55] and that was detected in the anagen hair follicles tested [56]. Several studies have indicated that VEGF, FGF2 and HGF were related to angiogenesis stimulating hair follicle proliferation [8; 57; 58; 59;60; 51; 62;63] and pro-angiogenic factor could promote anagen transition [64; 65]. Another previous study showed that Minoxidil, one of the pharmaceutical treatments approved for the treatment of alopecia, could promote hair growth by regulating VEGF expression in hair dermal papilla cells [66] and stimulated the transition to anagen [56]. AFMSC secrete MCP-1, IL-8, IL-6, EGF, SDF-1 and VEGF into the conditioned medium, which plays a vital role in angiogenesis [67]. Our data indicate that TGF-β and VEGF levels increased in the FAFI group, was remarkable, and this increase accelerated the transition to anagen phase and hair follicle growth, as stated in other studies. Furthermore, phagocytosis of cellular debris by M1 macrophages can promote production of TGF-β, one of the Th2 cytokines, while reducing expression TNF-α of the Th1 cytokines, reflecting their transition to the M2 phenotype [68]. In particular, the M1/M2 balance plays a critical role in determining the balance between fibrosis, inflammation/regeneration damage, together with the cytokine microenvironment. M2 polarization can be obtained in vitro with M-CSF, IL-4, IL-6, IL-10, IL-13, IL-33 and/or TGF-β, Arg+1 [69;70; 71; 72; 73; 74; 75; 76; 77; 78]. According to the concept of M1/M2 polarization IFN-γ, TNF-α and IL-1β are possible factors for the M1 macrophage activation [79]. A glycoprotein called CD68 is found on lysosomal membranes, specifically on the phagosomes in macrophages; its increased expression suggests improved phagocytosis [80] that is crucial in the process of tissue repair and wound healing. According to these results, our data showed that TGF-β was more prominent in the FAFI group so that induced M2 polarization. In addition, ≥ 10% of amniotic fluid concentrations were found to be related with the M1 to M2 phenotype switch in macrophages. In parallel with our study, Tan et al. [84; 85] reported that conversion of M1 to M2 in amnion epithelial cells content. M1-/M2-macrophages functionally correspond to Th1 and Th2, respectively [83]. In our study showed that the Th2 cytokine profile of the amniotic fluid was associated with M2 phenotype. Despite these limitations, hAF-conditioned media significantly altered the phenotype of macrophages toward an M2. Previous studies reported that AFSCs can provide protection by modulating immune function, especially by modulating macrophage recruitment and phenotype (by converting M1 macrophages to M2 macrophages) [84; 85; 86; 87]. According to researchers by Th2 cytokines increase the ornithine and urea production of arginase, which in turn increases macrophage arginine metabolism [88; 89].

In this study, we examined hAF treatment increased Arg+1 expression was observed around the hair follicles, and hair follicle was showed in the anagen phase especially in FAFI group. Our data indicated the expression of Arg+1, which is necessary in the regenerative hair follicle to demonstrate the activation of the M2 phenotype. Our study showed that CD68 was used marker to detect macrophages although other studies show that CD68 define as an M1 marker [90; 91]. We also observed that the expression of CD68 was observed in cnt group. These data demonstrate that CD68 associated with hair loss model.

HF degeneration is based on an anagen-related increase in macrophage numbers [19;92]. Previous study reported that skin-resident macrophages activated epithelial hair follicle stem cells that contribute to hair regeneration and induce anagen stage [44]. Chu et al. reported that by destroying M2 macrophages in mouse skin, hair regeneration was impaired but also it has been observed that there is an anagen onset when m2 infusion is performed so hair growth was observed in proportion to the number of M2 macrophages transplanted [23]. Our findings parallel with previous reports that the number of perifollicular Arg+1 as a used m2 marker increased during anagen and significantly decreased during catagen and reached the lowest levels during the telogen phase [43;93].

Our data showed that the second factor enabled the transition of M1 to M2 in hAF treated groups was that the amniotic fluid contains high exosome content (1.14e^10^±0.531e^10^). Indeed, in previous study showed that hAFSC exosomes contain HGF and TGF-β through characterized the extracellular vesicles secreted by AFSC [94]. Furthermore, hAFSC-exosomes treatment improved the regeneration levels of hair follicles, nerves and vessels [53].

Several studies have indicated that pre-adipocyte proliferation can be stimulated through angiogenesis [8], or cytokines such as PDGF, EGF, IGF1 and FGF [7; 95; 96], which activate dermal papilla cells to promote hair growth cycle and transition to anagen phase [6;7]. In our findings, thickness of adipocyte layer increased in the hAF-administered groups. Our data and previous findings [6; 7; 95; 96; 97; 98; 99; 100; 101; 102; 103] showed that thickness of dermal adipocyte layer increased during anagen phase but decreased during catagen and telogen phase. There is also evidence that immature dermal adipocytes activate HF stem cells to initiate the hair growth cycle [7]. In addition, the down growth of the HF during anagen may be facilitated by the presence of an adipocyte layer separating the reticular dermis from the underlying striated muscle, the panniculus carnosus.

In summary, we observed that the exosome content of the amniotic fluid and the contribution of growth factors enabled the conversion of M1 to M2 in the hair regerenation, stimulating the M2 polarization and increase in dermal adipocyte layer, which in turn stimulates the development of hair follicles. It is also thought that while gamma irradiation of the amniotic fluid increases the effectiveness of the product, possibly through protease inactivation, it also provides a safer product. These findings suggest that pooled and irradiated hAF may be involved in the clinical treatment of alopecia and in the pathobiology of hair follicle growth disorders. These findings are planned to be tested in a clinical trial. A new focus on irradiated amniotic fluid, which is safe, easily accessible and without provoking ethical concerns if applied in the clinic, promises to enrich not only translational hair research but also many other research fields as a model. In conclusion, our study revealed that hAF supports the hair follicle through cytokine, M2 and the extracellular vesicle. All these properties make them a promising source of FAFI for cell therapy and regenerative medicine.

## 4. Materials and Methods

### 4.1 Identification, Collection and Preparation of Amniotic Fluid Forms

hAF samples taken during cesarean section from 10 different donors who had previously approved amniotic fluid collection from the operating room of Acibadem Mehmet Ali Aydınlar University Atakent Hospital were transferred to the laboratory at 4°C. Pooling the amniotic fluid, which has low immunogenicity, from different donors further reduces the immunogenicity. Generally, it is recommended that relevant examination should be carried out before collecting hAF from healthy pregnant women, to avoid AF from patients with metabolic diseases. Researchers must also ensure that the hAF is free from viral such as hepatitis B and pathogenic bacterial infection [49]. Approximately 2 to 5 mL of hAF obtained via amniocentesis in the second trimester of pregnancy. Amniotic fluid was first passed through a 0.45 µl and then 0.22 µl filter in a laminar air flow cabin. An equal amount of pool was created by melting.

### 4.2 Preparation of Amniotic Fluid Forms (Fig. 4)

**Figure 4.**
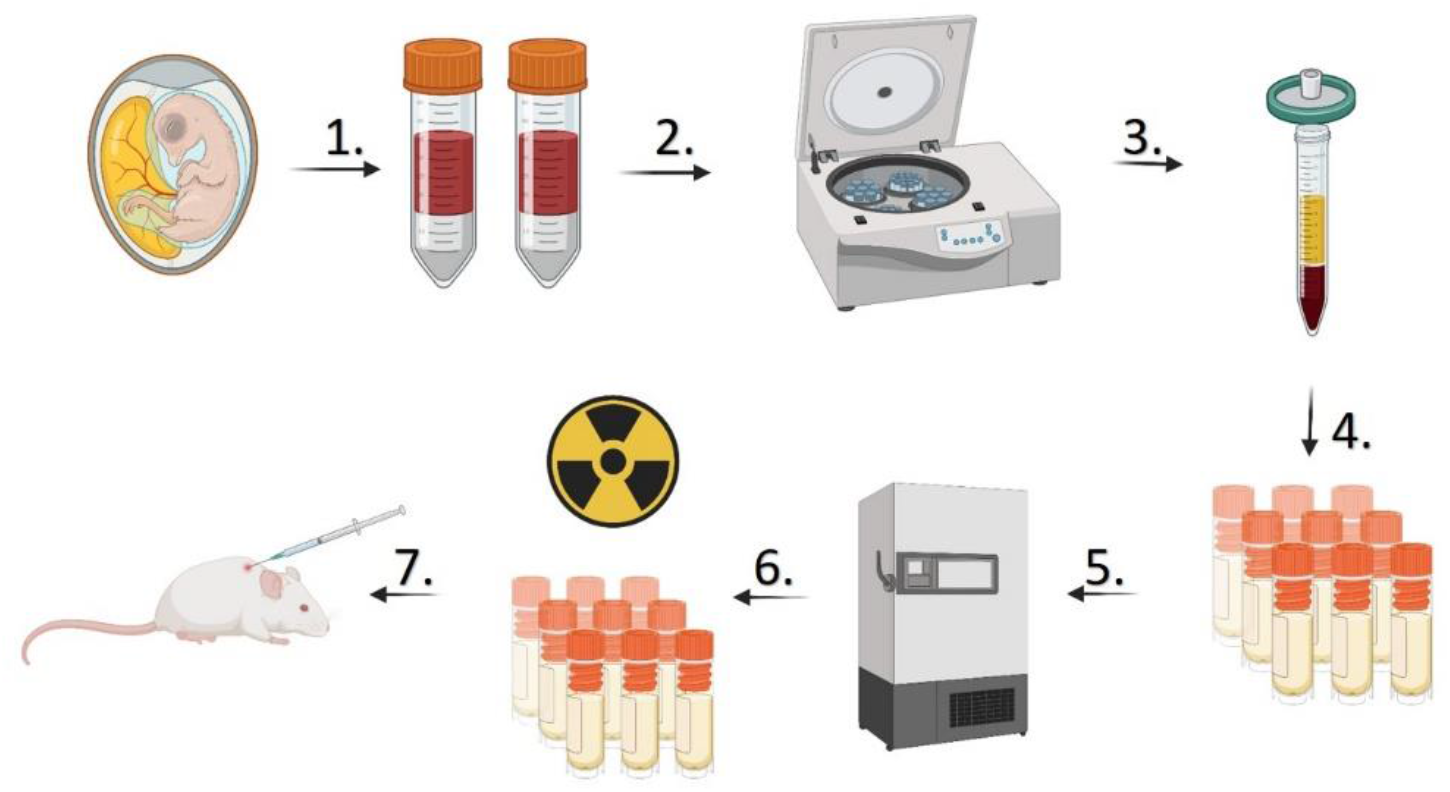
Schematic representation of hAF preparation. (1) Collecting amniotic fluid from at least three different sources. (2) Centrifuging amniotic fluids for the removal of macroparticles. (3) Staged filtration of centrifuged amniotic fluid. (4) Pooling of amniotic fluids from different sources together. (5) Freezing of the amniotic fluid at-80°C to −40°C. (6) Irradiation in frozen form between 25-35 Kgy. (7) hAF was applied subcutaneously.

#### FAF

Prepared amniotic fluid samples described above were dissolved into 2 ml 5 ml vials and then stored at −80 °C until the study period. (represents non-irradiated pool samples).

#### FAFI

FAF was exposed to 25 Kgray gamma radiation (irradiated FAF) in dry ice at −80°C. Before use, the contents of the vial were resuspended with 1cc sterile bidistilled water and used within 2 hours.

### 4.3 In vivo Animal and Experimental Design-Hair regeneration model

Our study Wistar albino female rats (6-8 weeks, weighing 250-300 g) were obtained with follow up with Acibadem Mehmet Ali Aydinlar University DEHAM (2019-32). The rats were anesthetized with 3% pentobarbital sodium (30 mg/kg) and dorsal skin hairs in rat were mechanical depilation. In experiment, rats were randomly assigned to 3 main groups (n = 7/group). (1) Cnt, The test group included a control group treated with normal saline; (2) Frozen amniotic fluid treated group; (3) Frozen irradiated amniotic fluid treated group (1 cc) subcutaneously to the on days 1, 3 and 5. Rats kept in temperature control and 12-hour light dark cycle and standart feeding in the nest. We considered all probable side effects including redness, swelling and rash during the experiment period. The rats were sacrificed and skin tissues were obtained on day 21. Photos were taken 21th days to record for macroscopic analyses. animals were anesthetized with subcutaneous injection of ketamine (100 mg/kg) and xylazine (5 mg/kg). They were horizontal biopsies were taken from the center of treatment areas.

After anesthesia and sacrificed according to the standart method. For histological analysis, dorsal skin samples were collected for each animal, samples were examined in a blinded fashion.

### 4.4. Amniotic fluid analysis: In the types of amniotic fluid used

#### 4.4.1 Cytokine ELISA Assay in Amniotic fluid

Protein standards for human growth factor (VEGF, EGF, PDGF-B, TGF-B, FGF-B, IL-1, IL-6) elisa strip (signosis) II (Cat# EA-1102). Add 200ul of Diluent buffer the wells of the first strip, and add 100ul of Diluent buffer to the wells of the rest strips according with the manufacturer’s protocol. The results were expressed as ng/l.

#### 4.4.2 Cytokine Bead Array (CBA) from Amniotic Fluid:Th1/Th2 cytokine ratios

Tumor necrosis factor alfa (TNF-α)/(IL)-10, TNF-α/IL-6, IFN-Ɣ/IL-6, interferon-alfa (IFN-α)/IL6, granulocyte macrophage stimulating factor (GMCSF)/IL-6, GMCSF/IL-10, GMCSF/IL-9 cytokine ratios of 1:2 diluted amniotic fluid were evaluated by the cytokine bead array (Miltenyi Biotec MACSPlex Cytokine 12 Kit, human; cat #130-099-169) in accordance with the manufacturer’s protocol. The experiment is repeated 3 times. Flow analysis were performed at Beckman coulter.

#### 4.4.3 Amniotic fluid induced macrophage differentiation

Peripheral blood mononuclear cell (PBMC) was isolated from whole blood collected in citrate tube by ficoll gradient method (GE Healthcare Bio-Sciences AB, Uppsala, Sweden). 500.000 PBMC /well was inoculated in 24 well plate in 1 ml. After 2 hours of incubation, suspended cells were dischared and adherent monocytes were observed. Dilutions were prepared with DMEM medium so that amniotic fluids were at a final concentration of 0%, 5%, 10%, and 20%. After 24 hours of incubation, the adheren monocytes were collected with trypsin and prepared for flow cytometry analysis. CD14-APC (cat#982506), CD86-Percpcy7 (cat#374209), CD206-FITC (cat#141703). Flow analysis were performed at Beckman coulter. Student t-test analysis were performed.

#### 4.4.4 Nanoparticle Tracking Analysis (NTA)

Microvesicle number was determined in amniotic fluid by using Te NANOSIGHT NS300 (Amesbury, UK). Samples were diluted with distilled water at a 1:10 ratio and transferred to NANOSIGHT cuvette as 1 ml. Measurements were performed at room temperature with 5 different 60-s video recording and were repeated 3 times.

### 4.5 Histopathological sampling analyzes

#### 4.5.1 Histochemical and Immunohistochemical stainings

Skin samples were fixed in 10% buffered neutral formol for 48 hours. The detected tissue samples were dehydrated by passing through alcohol series according to the histological tissue follow-up method, and after clearing in xylol, they were embedded in paraffin. Serial sections of 5 micron thickness were taken [50].

#### 4.5.2 Hematoxylin-Eosin (H&E) staining

Hair follicle parameters were stained with H&E (H3136 Sigma-Aldrich Hematoxylin and E4009 Sigma-Aldrich Eosin Y). Dermis-dermal adipocyte layer thickness and the numbers of hair follicles and anagen stages were counted in three areas (9 sections) from a biopsy on 3 random fields in 1 section using a high power microscope field by a histologist who was blinded to the codes.

#### 4.5.3 Masson-Trichrome staining

Collagen fibers were stained with Masson’s Trichrome staining (HT15-1KT, Sigma) kit to observe. Stained sections were evaluated for collagen determination in light microscopic images.

#### 4.5.4 Immunohistochemical staining

Sections of 5 µm thickness obtained from the selected paraffin blocks were taken to poly-L-Lysine coated slides and kept in an oven at 37 ^0^C for 1 night. Afterwards, deparaffinized sections were placed in xylene for 10 minutes and dehydrated by keeping them in 96% pure alcohol for 5 minutes. It was then washed in distilled water for 2 minutes. Following this procedure, in order to ensure antigen retrieval, it was heated in citrated buffer (ph: 6) solution at 98^0^C for 20 minutes and cooled at room temperature for 20 minutes in the same buffer. Afterwards, IHC staining was started. First, to eliminate endogenous peroxidase activity, it was blocked with 3% hydrogen peroxide, incubated for 20 minutes and washed with Phosphate Buffer Solution (PBS) for 5 minutes. Afterwards, protein block (Large Volume Ultra V Block, TA-125-UB®, Lab Vision Corporation, Fremont, CA, USA) was applied for 5 minutes. Following this procedure, before the sections were washed, the blocking solution was shaken off and the primary antibodies Arginase (Arg1 biocare cat#ACI3058) and CD68 (Dako cat#M0814) were applied. Afterwards, primary antibodies were washed in PBS for 5 minutes, secondary antibodies were dripped and incubated for 20 minutes. After washing for 5 minutes in PBS again, tertiary antibody was dropped and left incubated for 20 minutes. The 5 min PBS wash was repeated. Afterwards, chromogen was added to diaminobenzidine (DAB), incubated for 5-15 minutes and washed in distilled water. Tissues were counterstained in Mayer’s Hematoxylin for 1 min. Then it was washed in distilled water for 2-5 minutes and passed through alcohol. The air-dried preparations were placed in xylene, covered with entellan, and used for the specified measurements.

#### 4.5.5 Quantitative histomorphometry

Individual hair follicles in photomicrographs of H&E stained longitudinal sections of every rat. In photomicrographs in the same region (1300 m width) of the hair follicles were counted. In each group, the percentage of hair follicles that were in a particular anagen stage was calculated.

#### 4.5.6 Cell counting of Arg+1/CD68-polarized macrophages in in vivo skin analogs

In the sections, CD68 antibody staining evaluated as macrophage and Arg1 antibody staining evaluated as M2 rat macrophages of the cell density. In 9 different sections of each skin analog, 3 randomly selected were counted at 10× high-power fields. Cells showing distinct immunopositive reactions for CD68 and Arg+1 were counted per 0.2 mm^2^ from randomly selected areas in the FAF, FAFI and Cnt groups. The average total number of CD68 or Arg+1 positive cells quantified per high-power field of a skin section was used to express immune cell density. Results are given as the mean and standard deviation (SD).

#### Istatistical Analysis

The outcomes are shown as mean standard deviation (SD). One-way analysis of variance (ANOVA) was used to analyze the data. For all of our statistical analyses, we used SPSS version 23 the p-value.

## 5. Conclusions

As an alternative to cell therapy, especially gamma irridated pooled hAF may be beneficial in hair follicle regeneration treatments in the clinic. Moreover, hAF significantly enhanced their applications in experimental research and clinical practice.

## Abbreviations

AF: Amniotic fluid
AA: Alopecia Areata
AGA: Androgenic alopecia
Cnt: Control
DP: Dermal papilla
EGF: Epidermal growth factor
FGF-2: Fibroblast growth factor
GMCSF: Granulocyte macrophage stimulating factor
HFSC: Hair follicle stem cell
HF: Hair follicle
H&E: Hematoxylin-Eosin
hADSCs: Human amnion-derived stem cells
hAESCs: Human amniotic epithelial stem cells
hAMSCs: Human amniotic mesenchymal stem cells
hAFSC: Human amniotic fluid stem cell
hAF: Human amniotic fluid
IL: Interleukin
FAFI: Irradiated Frozen Amniotic fluid
IFN-γ: Interferon-gamma
IFN-α: Interferon-alfa
IGF-1: Insulin-like growth factor-1
FAF: Frozen Amniotic Fluid
M2: Macrophage2
M1: Macrophage 1
NTA: Nanoparticle Tracking Analysis
TGF-β: Transforming growth factor
TNF-α: Tumor necrosis factor
PDGF: Platelet-derived growth factor
PBMC: Peripheral blood mononuclear cell
VEG: Vascular endothelial growth factor

## 6. Patents

National (2022/009965) and international (PCT/TR2022/050661) patent numbers of this method have been received.

## Author Contributions

“Conceptualization, GTT. and EO.; methodology, GTT;EO.; software, GTT;EO.; validation, GTT;EO.; formal analysis, GTT. and EO..; investigation, GTT; BY;EG;DC.; resources, GTT; data curation, GTT.; writing—original draft preparation, GTT. EO..; writing—review and editing, GTT. and EO.; visualization, GTT. and EO.; supervision, GTT. and EO.; project administration GTT. and EO.; funding acquisition, EO.

## Institutional Review Board Statement

The ethical approval of this study was authorized by the Acibadem Mehmet Ali Aydinlar University Local Ethics Committee for Animal Experiments (ACU-HADYEK) with the decision number 2019/32 on the 12th of March 2019. All procedures in this study were performed in accordance with the 1964 Helsinki Declaration and its later amendments.

## Acknowledgments

In this section, you can acknowledge any support given which is not covered by the author contribution or funding sections. This may include administrative and technical support, or donations in kind (e.g., materials used for experiments).

